# Single-cell expression predicts neuron specific protein homeostasis networks

**DOI:** 10.1101/2023.03.14.532571

**Authors:** Sebastian Pechmann

## Abstract

The protein homeostasis network keeps proteins in their correct shapes and avoids unwanted protein aggregation. In turn, the accumulation of aberrantly misfolded proteins has been directly associated with the onset of aging-associated neurodegenerative diseases such as Alzheimer’s and Parkinson’s. However, a detailed and rational understanding of how protein homeostasis is achieved in health, and how it can be targeted for therapeutic intervention in diseases remains missing. Here, large-scale single-cell expression data from the Allen Brain Map is analyzed to investigate the transcription regulation of the core protein homeostasis network across the human brain. Remarkably, distinct expression profiles suggest specialized protein homeostasis networks with systematic adaptations in excitatory neurons, inhibitory neurons, and non-neuronal cells. Moreover, several chaperones and Ubiquitin ligases are found transcriptionally coregulated with genes important for synapse formation and maintenance, thus linking protein homeostasis to the regulation of neuronal function. Finally, evolutionary analyses highlight the conservation of an elevated interaction density in the chaperone network, suggesting that one of the most exciting aspects of chaperone action may yet be discovered in their collective action at the systems level. More generally, our work highlights the power of computational analyses for breaking down complexity and gaining complementary insights into fundamental biological problems.

## INTRODUCTION

Brain health critically depends on cellular protein homeostasis. In all cell types, a complex regulatory network of protein folding and quality control enzymes (Powers et al., 2009) such as molecular chaperones (Hartl et al., 2011) and Ubiquitin ligases (Zheng and Shabek, 2017) sustains the proteome in a folded and functional state. In turn, failure of protein homeostasis leads to the accumulation of misfolded and aggregation prone proteins (Hipp et al., 2019), and thus aging-related neurodegeneration (Chiti and Dobson, 2006).

Long studied primarily as a system of protein quality control (Neal et al., 2022), the protein homeostasis network readily extends in scope to serve as a central hub of regulating cell function (Rutherford and Zuker, 1994). One of the best demonstrations of this is found in neurons where synaptic plasticity can be controlled through the fine-tuning of protein levels between localized protein synthesis (Hafner et al., 2019) dependent on co-translational protein homeostasis (Pechmann et al., 2013), and regulated protein turn-over (Ross et al., 2021). Protein homeostasis thus directly couples selective protein quality control with regulation of neural activity (Giandomenico et al., 2022). However, a detailed understanding of protein homeostasis in general, and in neurons in particular, remains elusive.

Importantly, different tissues vary in the chaperones they express (Shemesh et al., 2021), as well as differentially predispose to protein homeostasis failure (Freer et al., 2016). Chaperones, especially co-chaperones that confer substrate specificity, dramatically increase in numbers with organism complexity (Draceni and Pechmann, 2019). Through tissue-specific expression (Shemesh et al., 2021), specialized protein homeostasis networks can both correspond to specific folding needs as well as achieve functional and regulatory fidelity. Yet, a main challenge of understanding protein homeostasis lies in its sheer complexity. This includes a large number of genes involved, a multitude of constraints on the expression of functional proteomes including for stoichiometry (Drew et al., 2021), function (Björkeroth et al., 2020), and solubility (Tartaglia et al., 2007), as well as strong systems-level synergies that achieve pervasive redundancy and resilience in folding (Rizzolo et al., 2017) and degradation (Breckel and Hochstrasser, 2021) networks. Because protein homeostasis demand and capacity are fine balanced (Draceni and Pechmann, 2019), comparative analyses of protein homeostasis in cells of specialized function such as neurons is poised to be particularly informative.

To this end, exactly how synapses form and are maintained within neurons also remains incompletely understood (Choquet and Triller, 2013; Südhof, 2018, 2021). From genomic and proteomic studies (Dieterich and Kreutz, 2016) a set of about 2000 genes likely important for the support of synapses has been curated (Pirooznia et al., 2012). Even less is known about the dependencies between synapses and protein homeostasis. For instance, only three E3 ligases could so far be linked to the regulation of pre-synaptic plasticity (Baccino-Calace et al., 2022).

Single cell transcriptomics now afford unprecedented insights into human cell types (Cao et al., 2019) including in the brain (Hodge et al., 2019; Özel et al., 2021). Remarkably, pronounced clusters of individual cells often suggest defined transcriptional cell identities akin to discrete and stable attractor states (Zhang and Wolynes, 2014). Because cell types often differ in the expression of thousands of genes, this has primarily motivated to understand cell differentiation (Dermadi et al., 2020; Maslovaa et al., 2020) as a problem of control, i.e. how these cell states are achieved, for instance through inference of the underlying transcription regulatory networks (Bendall et al., 2014). However, equally critical is to understand how the protein homeostasis network has tightly coevolved with the dynamic proteome demands to render these states functional.

Here, I present a large-scale comparative analysis on data from the Allen Brain Map of the transcription regulation of protein homeostasis at single-cell resolution across the human brain. Protein homeostasis networks of distinct and characteristic composition were found for excitatory neurons, inhibitory neurons, and non-neuronal cells. Moreover specific sets of chaperones and Ubiquitin ligases in these specialized networks were identified to be directly transcriptionally coregulated with the synaptome, thus suggesting possible functional links and candidate genes for future experiments. Finally, the high interconnectivity of the chaperone network more than individual interactions were found evolutionarily conserved, underlining the importance of understanding protein homeostasis at a systems level.

## RESULTS

The large-scale mapping of mRNA expression at single cell resolution has opened unprecedented opportunity to investigate shared and specialized characteristics of cell types. Here, single-cell expression data from the Allen Brain Map collected across multiple brain regions was analyzed to better understand whether neurons express specialized protein homeostasis networks that associate with synapse formation and maintenance. Specifically, by combining two recent studies on the systematic mapping of multiple cortical areas (Hodge et al., 2019), and the dorsal lateral geniculate nucleus (Bakken et al., 2021), expression profiles of 48642 neuronal and non-neuronal human cells partitioning into 131 clusters were obtained. Visualization in reduced dimensions supported the validity of merging these two studies, and highlighted a good separation between excitatory neurons, inhibitory neurons, and non-neuronal glia cells as overarching cell classes (Figure 1A).

**Figure 1.**
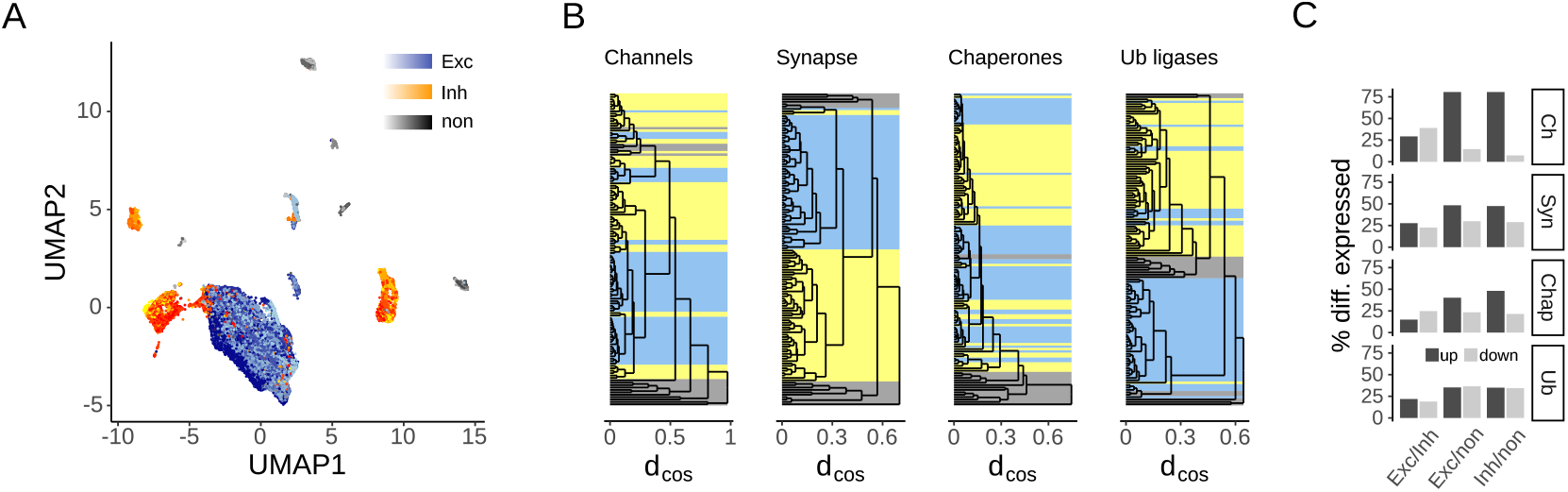
Expression of neuron-specific protein homeostasis networks in the brain. **(A)** UMAP plot of the single-nucleus mRNA expression profiles of 48642 cells in the human brain from the Allen Brain Map. Cells separated well into the cell classes of excitatory neurons (Exc), inhibitory neurons (Inh), and non-neuronal glia cells (non). **(B)** Comparison of the expression of protein homeostasis and synaptome genes. Dendrograms built from the hierarchical clustering based on cosine distance *d*_*cos*_ show similarities in the expression profiles of ion channels (n=42), synaptome genes (n=1746), chaperones (n=215), and Ubiquitin ligases (n=376). The corresponding cell classes are highlighted for excitatory neurons (yellow), inhibitory neurons (blue), and glia (grey). **(C)** Differential expression of protein homeostasis and synaptome genes between excitatory neurons, inhibitory neurons, and non-neuronal cells. For each pairwise comparison, indicated are the percentages of genes significantly up- and down-regulated.

As expected, the expression profiles of ion channels as a main characteristic of neurons strongly differentiated between excitatory neurons, inhibitory neurons, and non-neuronal glia cells (Figure 1B). The synaptome as the set of genes involved in synapse formation and maintenance (Pirooznia et al., 2012) is less defined, yet also readily partitioned cells by their class (Figure 1B). Within cell classes, for instance in comparison between excitatory neurons, differences were smaller as evident from shorter branch lengths in the cluster dendrograms (Figure 1B). After thus confirming that the differences in expression (Figure 1A) indeed traced back to genes especially important for neuronal identity, I asked whether such systematic differences also extended to the corresponding protein homeostasis networks. To test this, I compared the expression profiles of the core protein homeostasis network, namely the genes encoding the chaperone-mediated protein folding, and the Ubiquitin-dependent protein degradation systems (see *Methods*). Strikingly, the expression profiles of both chaperones and Ubiquitin enzymes were almost equally strongly predictive of the cell class, with a little more variability for the chaperones and an even stronger signal for the Ubiquitin enzymes (Figure 1B).

In direct comparison, ion channels were, naturally, strongly overexpressed in neurons (Figure 1C). Yet, the synaptome already painted a more varied picture. While many of its genes were overrepresented in neurons, a sizeable fraction was instead found enriched in glia cells (Figure 1C). The definition of the synaptome in its current form may habor potential for further context-dependent refinement. Interestingly, a large fraction of chaperones was also overexpressed in neurons, hinting at a prominent role of chaperone mediated protein homeostasis in neurons (Figure 1C). In constrast, some Ub enzymes were enriched in neurons and others in glia cells (Figure 1C). Moreover, several chaperone and Ubiquitin ligase genes were differentially expressed between excitatory and inhibitory neurons. Taken together, these observations established that excitatory and inhibitory neurons systematically express characteristic protein homeostasis networks of distinct composition.

How representative are these overall trends for individual cell clusters? Visualization of two-fold deviation from median expression highlighted extensive variability between clusters that was otherwise masked by summary statistics (Figure 2A). As reported previously (Shemesh et al., 2021), some chaperones were constituently expressed yet many others exhibited variable cell type specific expression. The same could be observed for Ubiquitin ligases (Figure 2A). Importantly, next to the systematic differential expression of protein homeostasis genes between cell classes, substantial variability was equally detected between different excitatory and inhibitory neurons respectively (Figure 2A).

**Figure 2.**
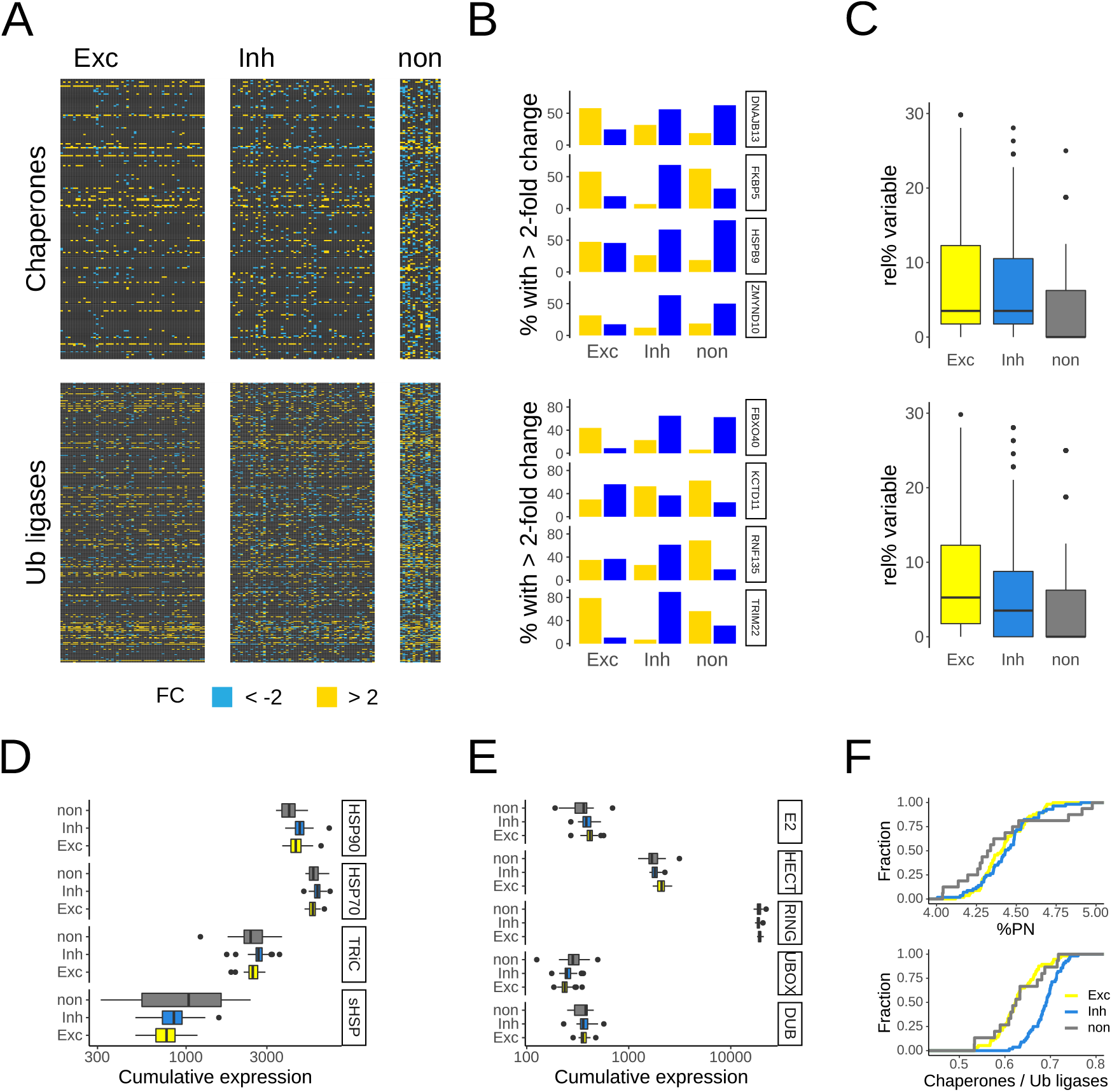
Variability and systematic adaptations in the expression of neuronal protein homeostasis. **(A)** Expression heatmap relative to the median of chaperone and Ubiquitin ligase mRNA levels; fold changes (FC) greater than two are shown. Columns denote individual cell clusters, and reported are average values for each cluster. Within the classes of excitatory neurons, inhibitory neurons, and non-neuronal glia, columns representing cell types are ordered following the hierarchical clustering based on their full genome expression similarity. **(B)** Percentages of exemplary chaperone and Ub ligase genes that were at least two-fold over- or under-expressed. Corresponding to the heatmap, substantial variability is observed for a large number of protein homeostasis genes even within classes of related cell types. **(C)** Distributions of the percentages of gene variability. For each chaperone or Ub ligase the percentage of cell clusters is shown within which the gene was identified as variably expressed. Chaperone genes were found particularly variable in excitatory neurons compared to their expression in the other cell classes. **(D)** Cumulative expression by enzyme family. To test if some variability of individual protein homeostasis genes is buffered at the systems level, cumulative expression levels for groups of genes that encode structurally, and thus often functionally, related proteins, are computed. Shifts in the distributions of cumulative expression levels suggest systematic adaptation in protein homeostasis network composition. **(E)** Distribution of the total expression of protein homeostasis gene expression relative to the full genome for all cell clusters and grouped by cell class. **(F)** Distribution of the ratio of chaperone to Ubiquitin ligase expression for all cell clusters and grouped by cell class. Inhibitory neurons express a much higher relative content of chaperones.

Several noteworthy examples stood out (Figure 2B). The HSP90 co-chaperone *ZMYND10* was preferentially found overexpressed in excitatory neurons and less frequently in inhibitory neurons and glia. In contrast, the HSP90 co-chaperone *FKBP5* was overexpressed in most excitatory neurons and glia, but not often in inhibitory neurons. The ER-localized HSP40 *DNAJB13* was found overexpressed only in excitatory neurons, while the sHSP *HSPB9* was overexpressed in half of the excitatory neurons, and depleted from the other half (Figure 2B). Similarly, the E3 Ubiquitin ligase *FBXO40* was overexpressed in excitatory neurons whereas *KCTD11* was depleted from most excitatory neurons but overexpressed in inhibitory neurons and glia. Another E3 ligase, *FNF135* was found to vary substantially between excitatory neurons while depleted from inhibotory neurons, and *TRIM22* was mostly depleted only in inhibotory neurons (Figure 2B). Remarkably, the same protein homeostasis genes were expressed more variably in neurons, especially excitatory neurons, than in non-neuronal cells (Figure 2C). These data suggest that individual cell type specific expression and pervasive variability are fundamental characteristics of protein homeostasis.

The presence or absence of individual protein homeostasis enzymes can be compensated by the expression of functionally redundant genes. To test whether the variable expression of some chaperones and Ubiquitin ligases was buffered at the systems level, I next looked at their cumulative expression. Chaperone families such as HSP90 or HSP70 type chaperones perform distinct tasks in the protein folding cycle (Hartl et al., 2011; Draceni and Pechmann, 2019), and thus comprise reasonable groupings. Importantly, the cumulative expression of chaperone families highlighted several interesting systematic adaptations. Specifically, the post-translationally acting stress chaperones from the HSP90 family were on average expressed more, from the sHsp family less in neurons compared to non-neuronal cells (Figure 2D). In contrast, the protein biogenesis HSP70s and the TRiC/CCT chaperonin subunits were systematically enriched in inhibitory neurons only (Figure 2D). Coordinating E2 ligases and deubiquitination enzymes (DUBs) perform unique tasks in the Ubiquitin-dependent degradation cycle, but groups of structurally related E3 ligases differ predominantly in their activation mechanisms rather than cellular function. Interestingly, E2 ligases were systematically more expressed in neurons compared to glia, UBOX-family ligases depleted in neurons, and HECT ligases enriched only in excitatory neurons (Figure 2E). The vast majority of E3s belongs to the family of RING ligases for which no pronounced difference was observed (Figure 2E).

Finally, the total expression of core protein homeostasis genes was found at a stable 4 *−* 5% of total expression irrespective of cell type or class (Figure 2F). However, the relative ratio between the cumulative expression of all chaperones, and all Ubiquitin enzymes respectively, revealed a strong and systematic shift of a much higher relative chaperone content in inhibitory neurons (Figure 2F). This observation traced back to the higher chaperone expression (Figure 2D) and lower Ubiquitin enzyme expression (Figure 2E) in inhibitory neurons. Thus, next to pronounced variability and cell-specific expression, these results revealed systematic adaptations in composition of the protein homeostasis networks in different cell classes.

Having established that neurons express protein homeostasis networks of distinct compositions, I next sought to test if these were linked to synapses as the their main function. Coexpression networks are a powerful tool for the discovery of coregulated genes that may be functionally coupled (Langfelder and Horvath, 2008). Here, selective coexpression networks were generated with the additional condition that only genes systematically overexpressed in target cells versus a control, for instance in neurons versus glia, were considered (see *Methods*). Owing to their central roles in cellular homeostasis, both chaperones and Ubiquitin enzymes had on average a higher degree of coexpression interactions than the rest of the genome (Figure 3A). Importantly, about 10% of the chaperones and Ubiquitin enzymes were both enriched in neurons and coregulated with the synatome (Figure 3B). Thus, a specific and select subset of protein homeostasis genes directly associated at the level of transcription with genes involved in synapse plasticity.

**Figure 3.**
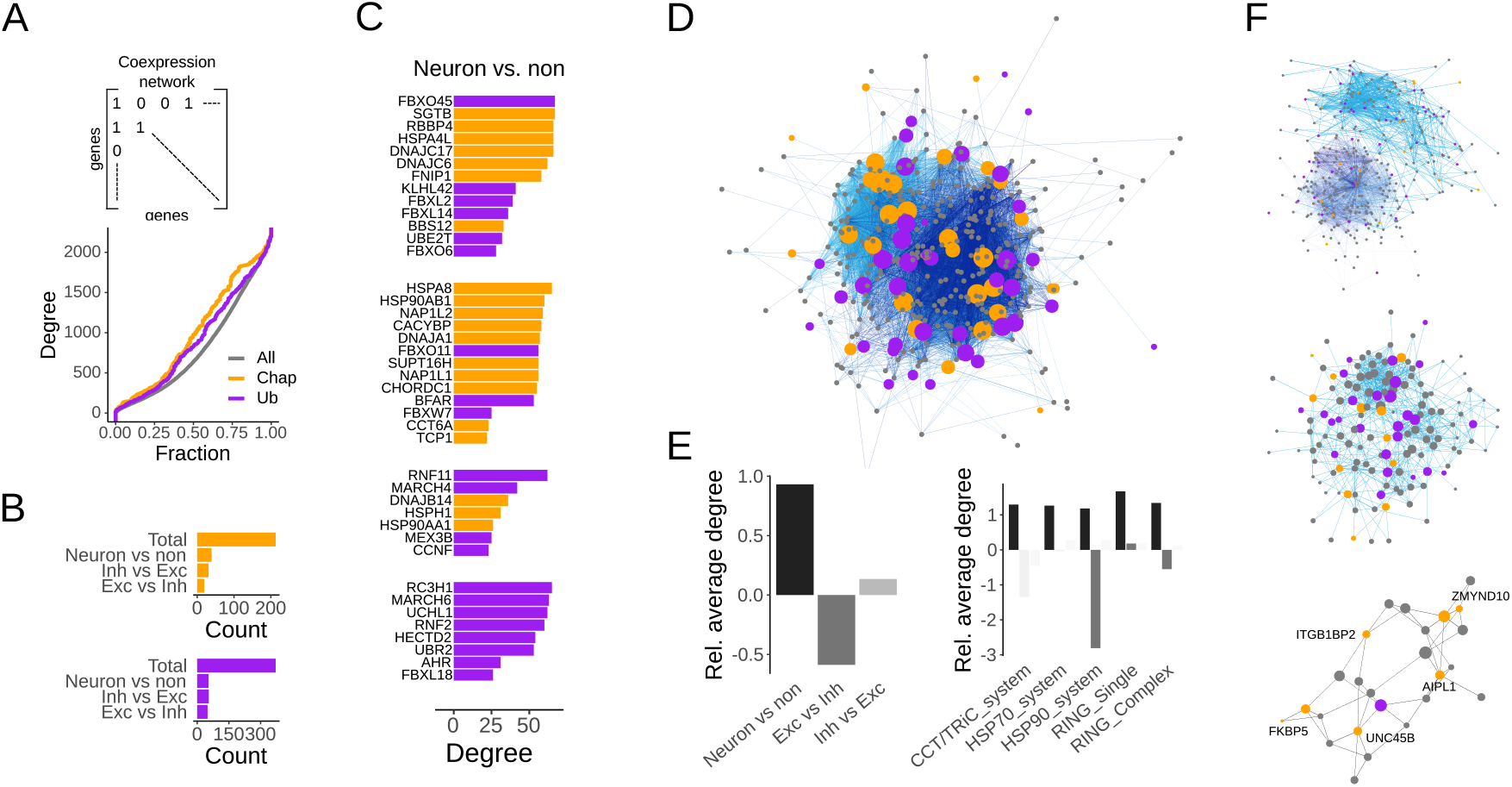
Coregulation of protein homeostasis and the synaptome. **(A)** Coexpression networks are built from coregulated gene pairs with highly correlated expression profiles. In the resulting networks, chaperones and Ubiquitin enzymes are characterized by above average degree distributions. **(B)** Selective coexpression networks are generated for all neurons, excitatory neurons, and inhibitory neurons. The number of chaperones and Ubiquitin enzymes interacting with the synaptome in each network are shown. **(C)** Protein homeostasis genes enriched in neurons and coregulated with the synaptome are shown together with their network degree and grouped by network modules. **(D)** Network representation of the coexpression network in neurons. The node size of protein homeostasis genes scales with their degree. **(E)** Relative network degree of protein homeostasis genes normalized by a randomization control and shown by cell class and corresponding enzyme family. **(F)** Selective coexpression network of excitatory neurons. Shown are the full network colored by modules (top), the module containing the most HSP90 co-chaperones (middle), and the subgraph containing only nodes connected to HSP90 co-chaperones (bottom).

The selective coexpression network of neurons compared to non-neuronal cells organized into several network modules that highlighted chaperones and Ubiquitin ligases coupled to the synaptome (Figure 3C). Strikingly, some of the thus identified genes were known to be directly involved in synapses, such as the E3 ligase *FBXO45* central to neuromuscular synaptogenesis. Others closely clustered, such as the HSP40s *DNAJC6* and *DNAJC17*, the HSP70 *HSPA4L*, and the HSP90 co-chaperone *SGTB*, all involved in conferring substrate specificity (Figure 3C). Another module contained three histone chaperones *NAP1L2, SUPT16H*, and *NAP1L1* as well as two subunits of the TRiCCCT chaperonin. While neither of these chaperones is neuron-specific, their context-specific coupling of chromatin regulation and translation regulation may be.

Overall, the resulting coexpression network between core protein homeostasis genes and the synaptome was dense and highly interconnected (Figure 3D). The relative connectivity of protein homeostasis genes, here defined as the number of coexpression interactions normalized by a randomized control (see *Methods*) was higher than expected in neurons (Figure 3E). This observation hinted at an elevated role of redundant and resilient protein homeostasis in neurons compared to non-neuronal cells. Furthermore, the direct comparison of excitatory and inhibitory neurons specified a slightly higher relative connectivity in inhibitory neurons and a clear reduction in the relative connectivity of protein homeostasis genes in excitatory neurons (Figure 3E). Breaking down these trends by enzyme family, the higher relative connectivity in neurons compared to glia was general. However, the reduction in relative connectivity observed in excitatory neurons traced down to the HSP90 system (Figure 2E). Moreover, by zooming in on the selective coexpression network in excitatory neurons relative to inhibiotory neurons, a network module containing several HSP90 co-chaperones could be identified (Figure 2F), thus suggesting indeed a functional link between them.

In conclusion, select subgroups of protein homeostasis genes were found not only systematically overexpressed in neurons but also transcriptionally coregulated with genes important for synapse formation and maintenance. Some of these genes were already known to be important for synapses, but the role of most has not been characterized in neurons. In addition, several systematic trends could be observed. HSP90 co-chaperones characterized by lower connectivity in excitatory neurons were consistently rediscovered in the same network module, likely because of intertwined functions. Naturally, the resolution of single-cell mRNA expression data is not sufficient to posit on mechanisms of protein interactions. However, the protein homeostasis genes identified should be some of the strongest candidates for future experiments into neuronal protein homeostasis given the current data.

Last, because evolutionary conservation is one of the strongest indications of functional importance, I sought to test whether the functional enrichment of protein homeostasis genes was evolutionarily conserved. To test this, species-specific coexpression networks were constructed for a smaller dataset of the dorsal lateral geniculate nucleus (Bakken et al., 2021) that contained expression data for human, macaque, and mouse (Figure 4A) (see *Methods*). As a control, I first looked at ribosomal genes that are known to be functionally coupled and conserved. Importantly, coexpression interactions between ribosomal genes were indeed strongly evolutionarily conserved compared to a randomized control (Figure 4B), suggesting informative power of such an analysis with the available data. However, interactions of chaperones and Ubiquitin enzymes were not more conserved than those of a random set of control genes (Figure 4B). This was not surprising given the promiscuous role of chaperones and Ubiquitin ligases that often interact with hundreds of client proteins, as well as a baseline signal reflecting the overall close relationship between the species.

**Figure 4.**
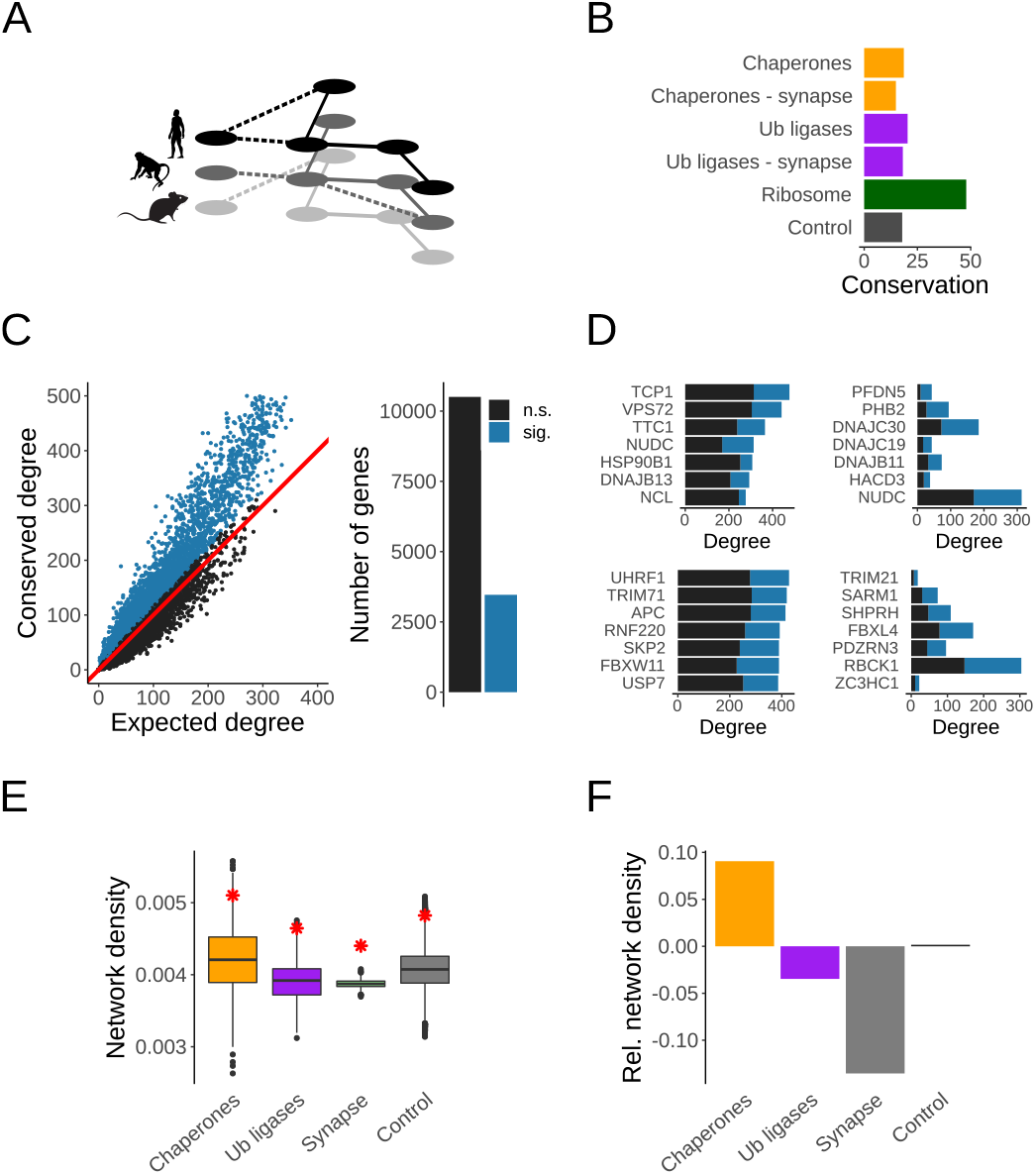
Evolutionary conservation of protein homeostasis coexpression networks in neurons. **(A)** Species-specific coexpression networks from human, macaque, and mouse single-cell expression data are used to test for evolutionary conservation. **(B)** Interactions between ribosomal genes as intrinsic control are well conserved, thus suggesting strong validity of this evolutionary analysis. Chaperone and Ubiquitin enzyme interactions are not more conserved than background. **(C)** Observed versus expected conservation of coexpression interactions. The interactions of only about a quarter of all genes are significantly conserved. **(D)** Protein homeostasis genes with the most conserved interactions (left) and the highest ratio of conserved to expected interactions (right). **(E)** Observed Network density of conserved interactions (red star) relative to the distributions of randomized controls for chaperone, Ubiquitin enzyme, synaptome, and control networks. **(F)** Relative network density obtained through normalization by a randomized control of the same network sizes affords the direct comparison and highlights a conserved higher network density in the chaperone network.

About a quarter of genes had more conserved coexpression interactions than expected by chance (Figure 4C). Of those, the chaperones with the highest number of conserved interactions included some of the most central chaperones such as the HSP90 *HSP90B1* and the TRiC/CCT subunit *TCP1* (Figure 4D). In contrast, the chaperones with the highest ratio of conserved to expected interactions were the prefolding subunit *PFDN5* as well as the three HSP40s *DNAJC30, DNAJC19*, and *DNAJB11* that are part of the triage of cotranslational protein folding. Similarly, the Ubiquitin ligases with the highest conserved degree included two E3s initially not linked to synapse biology, namely *UHRF1* important for chromatin regulation and *TRIM71* important for miRNAs, but also *APC* and *RNF220* that play important roles in Wnt-signaling, implicated in neurogensis. Of the Ubiqutin enzymes with the highest ratio of conserved to expected interactions, only *SARM1* is known to be important for protein degradation in axons while the others have not yet been implicated in neuron specific processes (Figure 4D).

Finally, protein homeostasis is characterized by strongly interconnected and redundant networks. The network density, here defined as the fraction of conserved coexpression interactions within a gene set, was found systematically higher than chance for all networks of chaperones, Ubiquitin enzymes, the synaptome and a randomized control (Figure 4E). Normalizing the observed network densities by the randomized controls to directly compare these networks of different sizes revealed a higher than expected relative network density of the chaperone network compared to the Ubiquitin network, and much higher than the network of the synaptome genes (Figure 4F). Interestingly, while both chaperones and Ubiquitin enzymes displayed on average a higher degree (Figure 3A), the conserved density was only higher in the chaperone network. It is well known that the chaperone network is especially modular, redundant, and dynamic, even more so than the Ubiquitin system that is more linear and hierarchic. The higher evolutionarily conserved relative network density across human, macaque and mouse transcriptomes highlighted the importance of systems level effects in addition to individual interactions especially for the chaperone system.

## DISCUSSION

This computational analysis of single-cell mRNA expression across the human brain has revealed protein homeostasis networks of distinct composition in excitatory neurons, inhibitory neurons, and non-neuronal cells. Remarkably, several protein homeostasis genes directly associated, through transcriptional coregulation, with genes involved in synapse formation and maintenance. While the thus identified chaperones and Ubiquitin ligases promise to be excellent candidate genes for future perturbation and characterization, beyond average variability in chaperone and Ubiqutin ligase expression as well as selection on systems level characteristics underlined the complexity and challenges of understanding protein homeostasis.

A major current limitation lies in mRNA expression levels as proxy for the inference of protein homeostasis networks, as mammalian cells utilize extensive translational regulation (Schwanhäusser et al., 2011; Cheng et al., 2016). Nonetheless, especially systematic and pronounced changes at the transcript level are strongly indicative of corresponding protein levels. The continued development of modern methods (Glauninger et al., 2022), especially for the quantification of protein abundances at the single cell resolution (Mair et al., 2020) will undoubtly further revolutionize insights and progress. Until then, there is tremendous merit in analyzing already available data within their limits.

One such limitation is an insufficient resolution to imply physical protein interactions between chaperones or Ubiquitin ligases and their clients from the transcriptional coregulation of their genes. To rationalize any regulatory interdependencies as well as how redundancies give rise to extraordinary resilience in protein homeostasis, interaction specificities in the network have to be better understood. Fortunately, many aspects of protein behavior can be increasingly inferred from their sequences and structures, such as their propensity for stress-granule formation (Boncella et al., 2020) or interaction with chaperones (Pechmann, 2020). Moreover, not all chaperone or Ubiquitin ligase interactions will involve functional regulation. Some of the most folding-challenged proteins such as acting and tubulin simply depend on specific chaperones for their folding needs (Gestaut et al., 2022; Rüssmann et al., 2012). In other cases, titration of shared chaperone or degrdation resources that prioritize the folding and quality control of some but not of other proteins may be much more exploited for systems level regulation than currently appreciated.

Despite tremendous achievements so far, further progress is also needed towards a rational understanding of synapse formation and maintenance (Südhof, 2021). Because neurons still cannot be easily cultured in the laboratory and thus subjected to all the technical advances of modern genomics, perturbation experiments are often cumbersome and limited to whole organism viable variants. Even more important become the use of already available technology and data together with integrated computational analyses that can help break down complexity and generate novel hypotheses. For instance the observed fundamental shift in the division of labor between chaperone-mediated protein folding and Ubiquitin-dependent protein degradation between excitatory and inhibitory neurons (Figure 2F) is remarkable because already an imbalance of excitatory and inhibitory neurons has contributed to aging (Wirak et al., 2022) and neurodegeneration (Maestuú et al., 2021).

Finally, the above average variability of chaperone and Ubiquitin ligase gene expression is pronounced and noteworthy. Gene expression in general is inherently stochastic (Raj and van Oudenaarden, 2008), as now quantifiable by single-cell genomics (Jariani et al., 2020). Moreover, some of that variability may be further amplified through translational heterogeneity (do Couto Bordignon and Pechmann, 2021). The suppression of expression noise is costly (Lestas et al., 2010), and some fluctuations are even functional, for instance in cell-fate decisions (Coomer et al., 2022) or during differentiation (Wada et al., 2018). However, a robust cell identity has to persevere in light of perturbations, for instance though robust control of transcription regulation (Mellis et al., 2021). Equally importantly, the expression of a functional proteome will have to be robustly supported by sufficient protein homeostasis capacity (Draceni and Pechmann, 2019). This is often a limitation in disease linked to protein misfolding (Hipp et al., 2019), but can also be a challenge in genome engineering or cellular reprogramming efforts, especially towards supporting synthetic non-native states (Zhang and Wolynes, 2014). The presented insight of cell specific protein homeostasis networks prepares for future mechanistic whole-cell models of protein homeostasis to address these challenges and better understand the role of protein homeostasis in health, aging, and neurodegeneration.

## METHODS

### Data and code availability

Project data and computer code generated for this project are available at https://www.github.com/pechmannlab/neuroPN.

### Data sources

Single-cell mRNA expression data was obtained from the Allen Brain Map (https://portal.brain-map.org/). Specifically, the processed and clustered human brain tissue single-nucleus (snRNA-seq) expression data from multiple cortical areas (Hodge et al., 2019) were analyzed together with a comparative analysis of single-cell expression data of the human, mouse, and macaque dorsal lateral geniculate nucleus, the brain region that relays visual information to the primary visual cortex (Bakken et al., 2021). Only cells with cluster assignments were analyzed, resulting in a set of 48642 cells in 131 clusters.

Gene annotations were obtained from literature. An encompassing set of around 2000 synaptome genes as an initial list of candidate genes linked to synapse formation, maintenance, and plasticity was obtained from the SynaptomeDB database (Pirooznia et al., 2012). A list of human genes encoding subunits of ion channels was taken from the ModelDB database (Mc-Dougal et al., 2017). A curated list of human chaperones (Brehme et al., 2014; Shemesh et al., 2021) was obtained from the Proteostasis Consortium (Elsasser et al., 2022), and a list of genes in the human ubiqutin-dependent degradation system was compiled from the hUbiquitome (Du et al., 2011) and UbiNet2.0 (Li et al., 2021) databases.

### Data analysis

All analyses were limited to genes whose annotations uniquely mapped to corresponding protein sequences in the non-redundant Uniprot reference proteomes (The UniProt Consortium, 2021). Expression levels were normalized to counts per million. Differentially expressed and variable genes were detected as implemented in Seurat (Butler et al., 2018). Dimensionality reduction of cell expression profiles was performed with UMAP (Becht et al., 2019).

### Coexpression networks

To circumvent the problem of missing values in single cell expression data, the data was transformed to a vector of pairwise comparisons that by themselves were more robust (Iacono et al., 2019). Specifically, the per gene expression profile across *n* cell clusters was replaced by a vector of *n ∗* (*n −* 1) fold changes from pairwise differential expression tests. Correlation coefficients between gene pairs were then used to construct coexpression networks (Langfelder and Horvath, 2008). Thresholds to call coexpression interactions were arbitrary and reflected a trade-off between stringency and sufficient data points for further analysis; in the full data the top 5%, and in the smaller species-specific coexpression networks the top 20% of correlating gene pairs were called as interacting. Furthermore, to construct selective coexpression networks, only genes that were systematically overexpressed by at least 20% in the target cells versus a control, for instance in all neurons versus non-neuronal cells, were considered. Last, to increase confidence and limit networks to genes that may partake in feedback interactions, all motifs of size *n* = 3 within coexpression networks were enumerated as previously described (Sahoo and Pechmann, 2022). Only genes contained in the motifs, i.e. with multiple systematic interactions, were subsequently used to construct final networks. Network modules were detected with the Python networkx library.

### Network evolution

Ortholog assignments for the reference proteomes without isoforms (The UniProt Consortium, 2021) between human, macaque, and mouse were recomputed with OMA (Altenhoff et al., 2019). Comparative expression data for these three species was available from (Bakken et al., 2021). Coexpression networks were generated as described above, and interactions between orthologous gene-pairs that were present in all species-specific networks were considered as conserved. To understand any selective forces on the conservation of coexpression interactions, null-models were generated through degree-preserving randomizations of the species networks with the R package BiRewire (Iorio et al., 2016). Relative connectivity was defined as the observed network degree relative to the corresponding averages resulting from randomized networks of the same sizes.

## ACKNOWLEDGMENTS

This research was carried out while caring for a family member.

## DISCLOSURE DECLARATION

The author declares no competing interests.

## Notes

https://www.github.com/pechmannlab/neuroPN

### Competing Interest Statement

The authors have declared no competing interest.

